# Gretl - Variation GRaph Evaluation TooLkit

**DOI:** 10.1101/2024.03.04.580974

**Authors:** Sebastian Vorbrugg, Ilja Bezrukov, Zhigui Bao, Detlef Weigel

## Abstract

Motivation: As genome graphs are powerful data structures for representing the genetic diversity within populations, they can help identify genomic variations that traditional linear references miss, but their complexity and size makes the analysis of genome graphs challenging. We sought to develop a genome graph analysis tool that helps these analyses to become more accessible by addressing the limitations of existing tools. Specifically, we improve scalability and user-friendliness, and we provide many new statistics for graph evaluation.

Results: We developed an efficient, comprehensive, and integrated tool, *gretl*, to analyse genome graphs and gain insights into their structure and composition by providing a wide range of statistics. *gretl* can be utilised to evaluate different graphs, compare the output of graph construction pipelines with different parameters, as well as perform an in-depth analysis of individual graphs, including sample-specific analysis. With the assistance of *gretl*, novel patterns of genetic variation and potential regions of interest can be identified, for later, more detailed inspection. We demonstrate that *gretl* outperforms other tools in terms of speed, particularly for larger genome graphs.

Availability and implementation: *gretl* is implemented in Rust. Commented source code is available under MIT licence at https://github.com/MoinSebi/gretl. Examples of how to run *gretl* are provided in the documentation. Several Jupyter notebooks are part of the repository and can help visualise *gretl* results.

## I. Introduction

Advances in short-read based resequencing have greatly improved our understanding of genomic variation in many different species (1001 Genomes Consortium, 2016; Peter *et al*., 2018; 1000 Genomes Project Consortium, 2015). More recently, long reads have made it possible to assemble complete genomes with remarkable speed and precision. Moving from variant inventories to complete genomes facilitates much more comprehensive analysis and genome-wide comparison between samples. As an example, in the plant *A. thaliana*, the level of detail provided by (nearly) complete genomes has already led to new insights into conservation of synteny and to much more accurate description of single nucleotide polymorphisms (SNPs), copy number variants (CNVs), and structural rearrangements (Jiao and Schneeberger, 2020; Goel *et al*., 2019).

To mitigate the biases associated with a single reference genome, pan-genomes built from diverse sample collections are being created from increasingly complex genomes (Liao *et al*., 2023; Shang *et al*., 2022). A crucial tool for efficient storage and comprehensive analysis of genetic variations within diverse and intricate genomic regions is the variation graph, which condenses similar sequences into nodes and captures variations in a reference-free manner. Graph shape and structure depend on the choice of construction method and parameter set, requiring tuning and adjustment based on the genome complexity (Crysnanto *et al*., 2022) and the research question, highlighting the need for a comprehensive evaluation tool.

Genome graphs are typically stored in GFA (Graphical Fragment Assembly) format, a standardised data format, which is also the main input for *gretl*, the tool introduced here. Nodes in the graph represent DNA segments, connected by edges and each node has a unique identifier and associated DNA sequence. GFA can store additional information like allele frequency, quality scores, or annotations, if needed. The format ensures interoperability among software tools, facilitating collaboration and analysis development. *gretl* fully supports GFAv1 files (http://gfa-spec.github.io/GFA-spec/GFA1.html), ensuring interoperability across a wide range of graph tools. Adopting GFAv2 is an option for the future, as more upstream graph tools migrate to GFAv2.

Several tools for genome graph analysis are available and being actively developed, including *odgi* (Guarracino *et al*., 2022), *vg* (Garrison *et al*., 2018) and *gfastats* (Formenti *et al*., 2022). While *odgi* and *vg* offer powerful platforms for modifying and analysing genome graphs, they lack certain statistical features crucial for large-scale graph analysis. Although *gfastats* is designed for statistics, its primary focus lies in assembly graphs, which have, in comparison with whole-genome graphs, distinct characteristics. While it does provide several useful statistics for genome graphs, its main function remains an overall toolkit for modifying GFA files and delivering high quality single individual genomes.

One of the primary motivations behind our work was to provide a fast and efficient tool for the initial evaluation of newly constructed graphs. Building genome graphs is a complex process, and one often needs to rapidly assess their quality. *gretl* aims to address this need by offering an all-in-one tool that evaluates graph structure and composition (Figure 1A).

**Figure 1.**
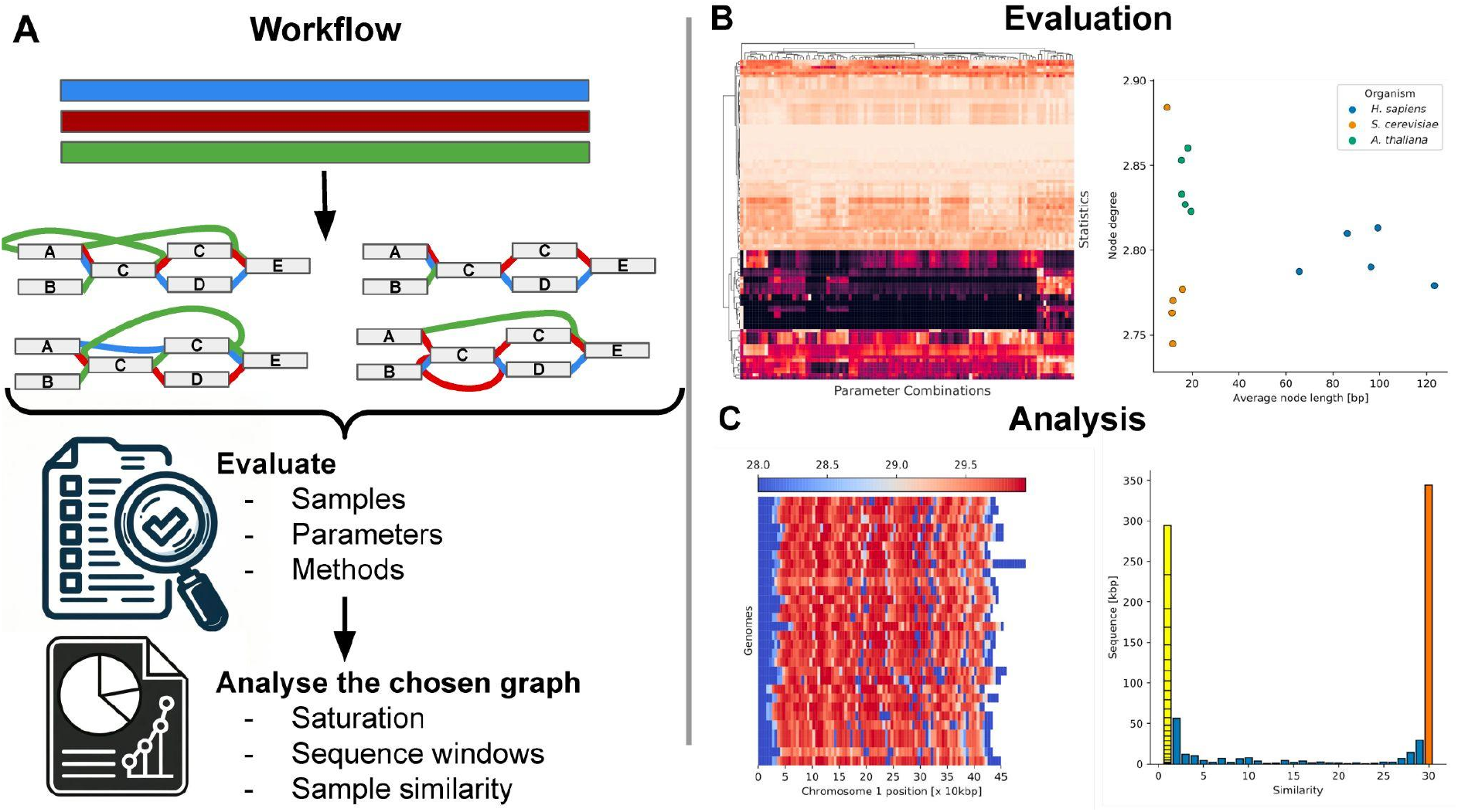
Gretl overview. (A) Genome graph construction workflow: Genome graph properties are influenced by various factors, including parameter selection, sample curation, and methodology, all of which impact the layout and structure of the resulting genome graph. For evaluation purposes, multiple graphs may be simultaneously generated and must be evaluated to identify an optimal representation for a specific task. The selected graph can then be analysed with gretl. (B) Visualization of gretl output: Left, scatter plot depicting two selected statistics across various graphs, facilitating comparisons between different species. Right, graphs can be clustered based on multiple statistics, grouping similar species or construction parameters (shown here). (C) In-depth analysis of a selected genome graph (example yeast): Left, pan-genomic analysis of the genome graph. Sequences found only in a single sample are separated and each block represents one path of the graph. Right, path-centric sliding window analysis of the S. cerevisiae genome graph, highlighting regions of high similarity.

With *gretl*, researchers can assess the overall integrity of the graph and identify potential areas that require further investigation or refinement. By automating the analysis process, we eliminate the need for manual evaluations. Moreover, one can generate statistics on genome-growth, pan-genome distribution and/or for specific paths (Figure S1, S6). This preliminary information can guide subsequent analyses and describe properties of different species pangenomes, providing a solid foundation for further investigations.

## II. Results

*gretl* offers valuable insights into genome graphs constructed using PGGB (Garrison *et al*., 2023) and Minigraph-Cactus (Hickey *et al*., 2023), as well as other graphs in GFA format. *gretl* provides several subcommands that offer comprehensive graph analysis, covering aspects such as graph complexity, interconnectedness, and node degree. We provide Jupyter notebooks that can be used for post-processing and visualisation of the output, allowing further exploration and interpretation of the results.

To illustrate the capabilities of *gretl* for graph evaluation, we used pan-genome graphs constructed from chromosomes of *H. sapiens* (n=48 for Chr 14, 18, 19, 21, 22), *S. cerevisiae* (n=30 for Chr 1, 3, 5, 9, 10) and *A. thaliana* (n=67 for Chr 1-5). Details about the datasets (Liao *et al*., 2023; O’Donnell *et al*., 2023; Wlodzimierz *et al*., 2023) and graph construction are given in Supplementary Data.

Firstly, the provided statistics enable evaluation of graphs built with different parameters from the same dataset (Figure 1B). Statistics generated by *gretl* (Figure S1, S6) can guide subsequent analyses and describe the properties of different species pangenomes.

Additionally, these values facilitate the evaluation and comparison of graphs from different methods or organisms, enabling insights into the complexity and structure of the genome (Table S1). It is important to note that some of these statistics may exhibit similar behaviour and display high correlation due to their interconnected nature (Figure S5). Researchers can explore the impact of varying parameters during graph construction on the same dataset or analyse different species by comparing or clustering their genome graphs by statistical features (Figure 1B).

In general, *gretl* offers more comprehensive information about the graph than other tools. The tabular output format provides an easy overview as well as seamless integration with scripting languages such as R and Python for post-processing.

Secondly, *gretl* facilitates in-depth comparison of specific graphs using a wide range of metrics. This analysis can be performed at both the graph level and the path level, providing researchers with comprehensive insights. At the graph level, various metrics and statistics can be explored to identify regions of interest, which can be further investigated in subsequent studies (Figure 1C). Sliding window analyses on sequence or node level give powerful insight into local complexity or distant sequence similarities (Figure S7).

The path-centric analyses allow for computation of independent statistics and metrics for specific paths within the graph. This approach enables comparisons between different samples or populations, helping in the identification of path-specific differences within the pan-genome (Figure 1C). Furthermore, it enables the identification of samples that display an isolated or extreme representation in the graph (Figure S3). By carefully examining the paths within the graph, researchers can uncover structural patterns, variations, and potential functional significance embedded within the genome. This comprehensive analysis of paths should contribute to a deeper understanding of the genome’s complexities and provides valuable insights for further research.

During testing on a 3.2 GHz AMD Epyc 64 core machine using the HPRC graph (Liao *et al*., 2023), which consisted of 48 samples (96 haplotypes, 1882 samples) with 4.15 million nodes and 5.79 million edges, our evaluation tool demonstrated a level of performance that should greatly encourage its adoption for any genome graph construction workflow. It computed simple summary statistics twice as fast as other approaches (^∼^8 min), utilising 4.3 GB of memory (Figure S2).

## III. Discussion

We contribute *gretl*, a fast, efficient, and user-friendly stand-alone tool for generating a wide range of statistics and insights into the structure and composition of genome graphs, complemented with a set of user-friendly Jupyter notebooks for downstream analyses. *gretl* generates 108 different metrics for a single variation graph.

It is important to note that the quality of the genome assemblies used to generate the genome graph can significantly affect the accuracy and completeness of the generated metrics and subsequent downstream analyses. As such, it is essential to carefully evaluate and validate assembly quality before using *gretl*. In our experience, the building of graphs from complex genomes such as those of plants is highly affected by parameter choice.

While *gretl* can process any graph which adheres to the GFAv1 specification, it has specific requirements for node identifiers to produce consistent results. Specifically, the node IDs should be numerical, consecutive, and sorted in a manner that follows the pan-genomic (consensus) order. This ordering of the graph can be achieved effectively using the ‘odgi sort’ functionality during the preprocessing stage (Guarracino *et al*., 2022; Heumos *et al*., 2023). *gretl* relies on sorted genome graphs for its time-efficient processing and several key statistics, such as “Edge Jump” and “Path jumps”, which are based on a graph layout.

*gretl* is unique in that it provides both graph-based and path-based statistics, allowing users to gain insights into both the overall structure of the genome graph and the specific paths/samples through the graph that correspond to genetic variation. Finally, *gretl* is designed to be modular and extensible, allowing for the future addition of new features and statistics.

## Data and code availability

Source code is available under MIT licence at https://github.com/MoinSebi/gretl together with Jupyter notebooks.

## Acknowledgements

We thank Christian Kubica and Simon Heumos for discussion.

## Supplementary data

Supplementary data are available at *Bioinformatics* online.

