## Supplementary Information for "Gretl - Variation GRaph Evaluation TooLkit"

### **Gretl** - Variation **GR**aph **E**valuation **T**oo**L**kit

### 1. Implementation

For optimization purposes, ***gretl*** has been implemented in the Rust programming language. It incorporates several Rust crates to enhance performance and enable multithreading. Furthermore, a new GFA format reader has been integrated as a library. In addition to handling the GFAv1 format utilized by ***gretl***, this reader is capable of handling GFAv2 as well and can be used by other tools.

### 2. Datasets

The datasets used in this study were sourced from public resources and previous publications. *A. thaliana* genomes were obtained from (Wlodzimierz *et al.*, 2023), while *S. cerevisiae* genomes were collected and sequenced as described in the publication by (O'Donnell *et al.,* 2023) . The human graph evaluated in the figure was directly downloaded from https://github.com/human-pangenomics/hpp_pangenome_resources and is a part of the dataset introduced by the draft human pan-genome (Liao *et al.*, 2023).

### 3. Building a pan genome

Except for human data, graphs were constructed using the PGGB pipeline (Garrison *et al.*, 2023) for each chromosome individually. As previously mentioned, only graphs constructed from human assemblies were downloaded from https://github.com/human-pangenomics/hpp_pangenome_resources. The PGGB pipeline was executed with default parameters for the species comparison experiment.
Parameter comparisons were conducted using *S. cerevisiae* genomes with various parameter combinations.
The parameters we modified included -s (2k, 5k, 10k), -p (80, 90, 95), -k (19, 31), -n (15, 30, 60), and -P (asm5, asm10). The ***pggb*** workflow was executed with ***wfmash*** (v0.10.2-2-gb310bd1), ***seqwish*** (v0.7.8-3-gd9e7ab5), ***odgi*** (v0.8.2-92-gbfae0b3), and ***smoothxg*** (v0.6.8-31-g06bbf35).
*A. thaliana* graphs were constructed with default parameters.

### 4. Pan-genome classification

We utilised the characteristics of the graph to classify different levels of relatedness for *S. cerevisiae* genomes. We annotated nodes as core, soft, and private using the following approach. Nodes which were present in all accessions, were labelled as core. In contrast, nodes that were only traversed by one accession are private, while everything between these values (>1 and <30) were classified as soft (shell) nodes.

### 5. Table of all reported features (*gretl* stats)

A table of all reported features can be found in the github repository: https://github.com/MoinSebi/gretl/paper. Order of the statistics might change over time, but the linked table will remain stable to mirror the features represented in this publication.

| Organism | Chr | #Sequences | #Samples | #Nodes [x1000] | #Edges [x1000] | Average node size [bp] |
|---|---|---|---|---|---|---|
| ***Homo sapiens*** | **14** | 1882 | 48 | 4,154.6 | 5790.4 | 65.68 |
| | **18** | 1270 | 48 | 2,832.4 | 3979.5 | 86.17 |
| | **19** | 1072 | 48 | 3,021.1 | 4214.6 | 96.24 |
| | **21** | 3029 | 48 | 2,760.5 | 3882.9 | 99.19 |
| | **22** | 1757 | 48 | 3,759.7 | 5224.4 | 123.37 |
| ***Saccharomyces cerevisiae**** | **I** | 30 | 30 | 52.3 | 75.8 | 9.27 |
| | **III** | 30 | 30 | 52.7 | 72.9 | 11.41 |
| | **V** | 30 | 30 | 125.2 | 171.4 | 15.90 |
| | **IX** | 30 | 30 | 67.3 | 93.2 | 11.84 |
| | **X** | 30 | 30 | 73.6 | 101.0 | 11.71 |
| ***Arabidopsis thaliana***** | **1** | 67 | 67 | 6,891.3 | 9,740.7 | 17.00 |
| | **2** | 67 | 67 | 4,926,7 | 6,978.9 | 15.54 |
| | **3** | 67 | 67 | 5,972.6 | 8,520.0 | 15.43 |
| | **4** | 67 | 67 | 4,747.1 | 6,788.9 | 18.16 |
| | **5** | 67 | 67 | 5,656.715 | 7,984.3 | 19.58 |

**Supplementary Table 1**: Information on the genome graphs used in this study. *Parameter set -p 90, -n 30, -s 5000, -k 31 **Default parameters

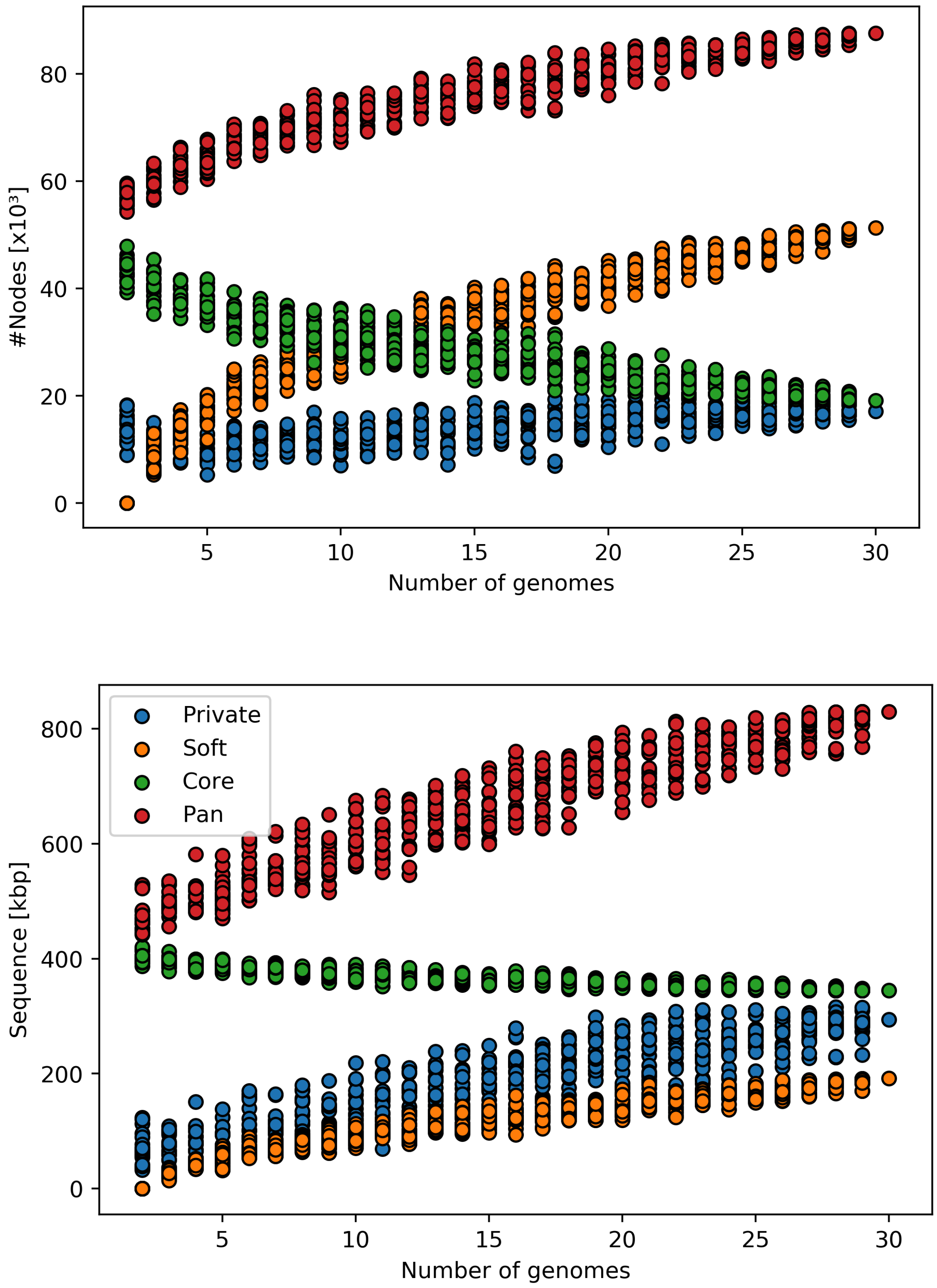


**Supplementary Figure 1**: Saturation analysis using a 20x bootstrap approach based on the *S. cerevisiae* genome graph for chromosome IX.
Top figure represents the amount of sequence in each category, bottom figure displays the number of nodes**.** Classification has been applied as described in Section 4. The pan-genome does not seem to saturate, which is mainly pushed by new variations being added with additional genomes.

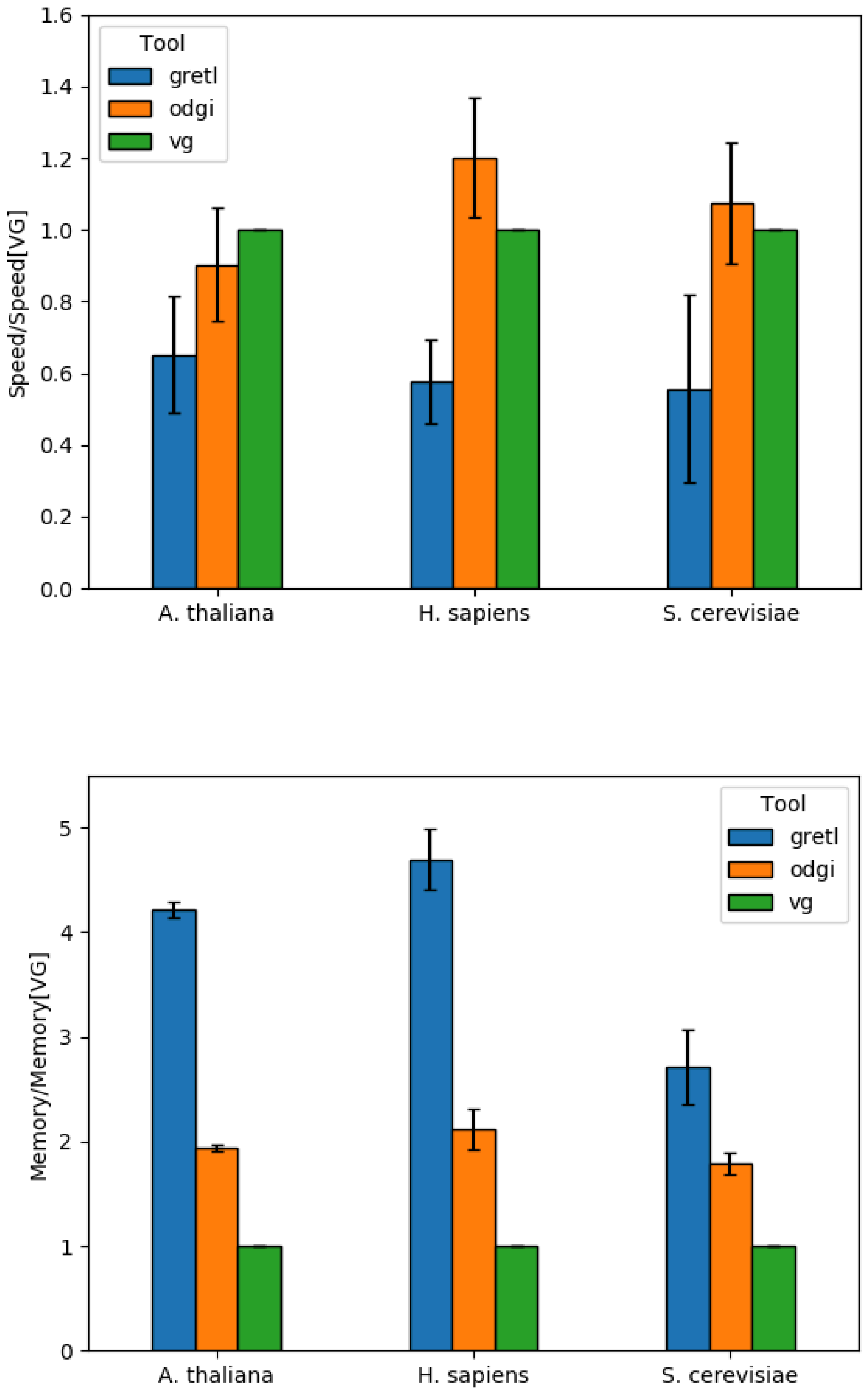


**Supplementary Figure 2**: Run-time and memory benchmarking for ***gretl***, ***vg stats*** and ***odgi stats*** in relation to ***vg stats***.

For each species five graphs were used (s. Table S1). In general, ***gretl*** is faster but requires more memory than other tools.

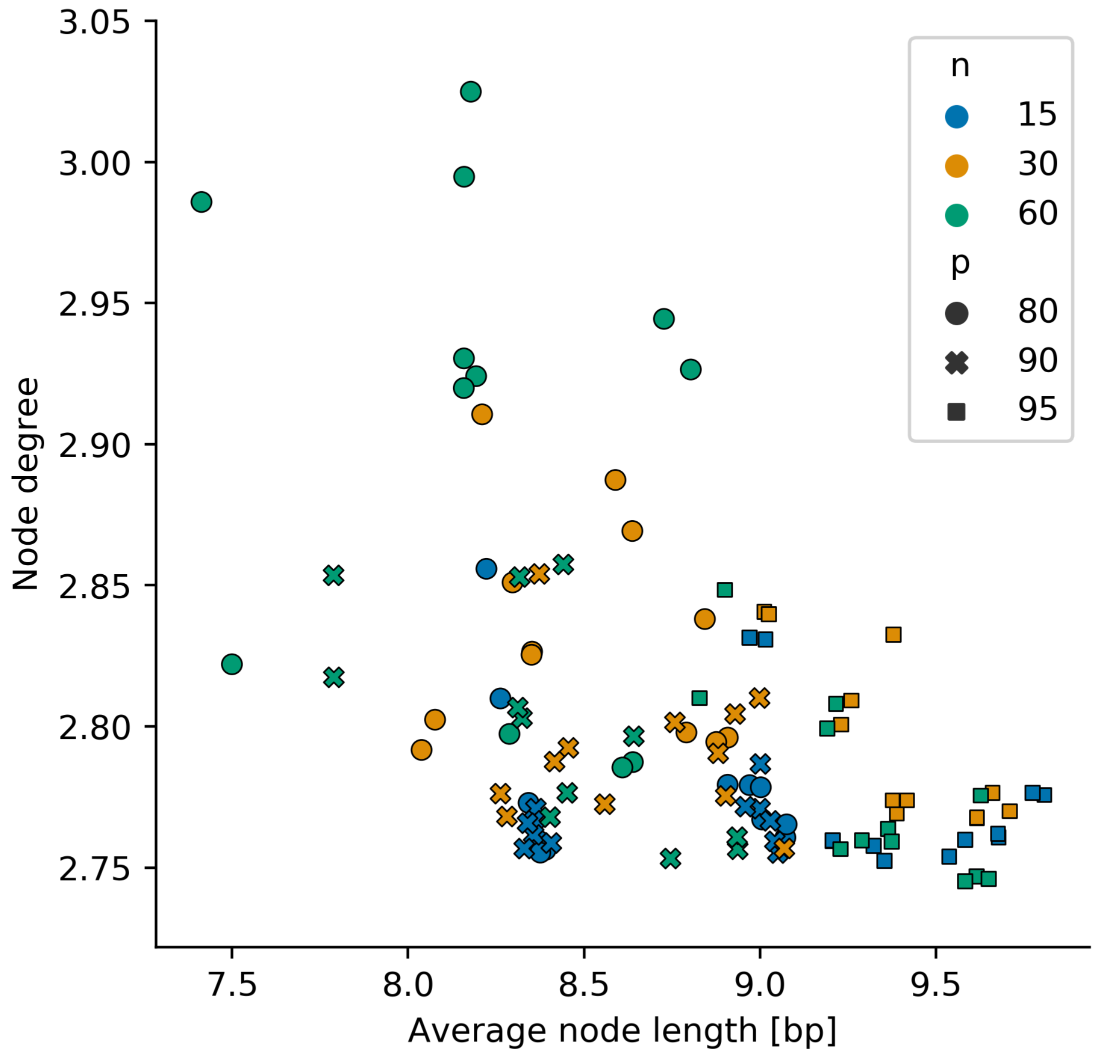


**Supplementary Figure 3**: Scatterplot highlighting the node degree and the total number of nodes with a single base pair for all graphs built with different combinations of parameters. The different colours highlight the “percent identity” in the ***wfmash*** step (-p), the different shapes the (secondary) n-mappings (-n) of ***pggb***. Graphs are based on 30 *S. cerevisiae* genomes of chromosome IX. Graphs built with different “p” can easily be distinguished and this parameter seems to have a strong influence on the graph.

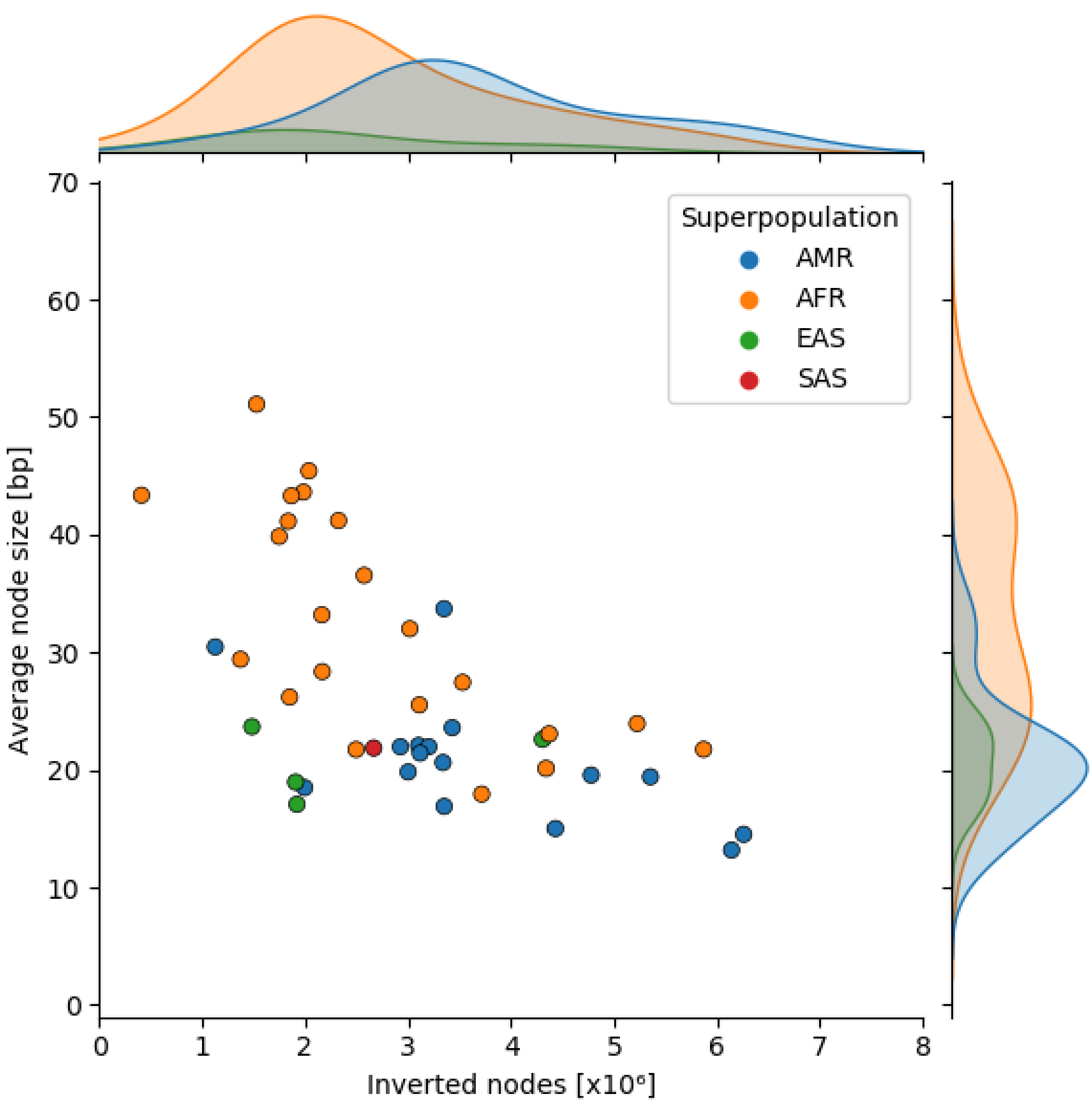


**Supplementary Figure 4**: Scatterplot highlighting the average node size and the number of inverted nodes [x10$^6$] of each path in the *H. sapiens* graph.
The path names are annotated to their respective data point. The graph was built with chromosome 1 of 30 *A. thaliana* genomes using the ***pggb*** pipeline (-p 90, -n 30, -s 10000, -k 31). Genomes divergent in sequence context can be easily identified using graph statistics.

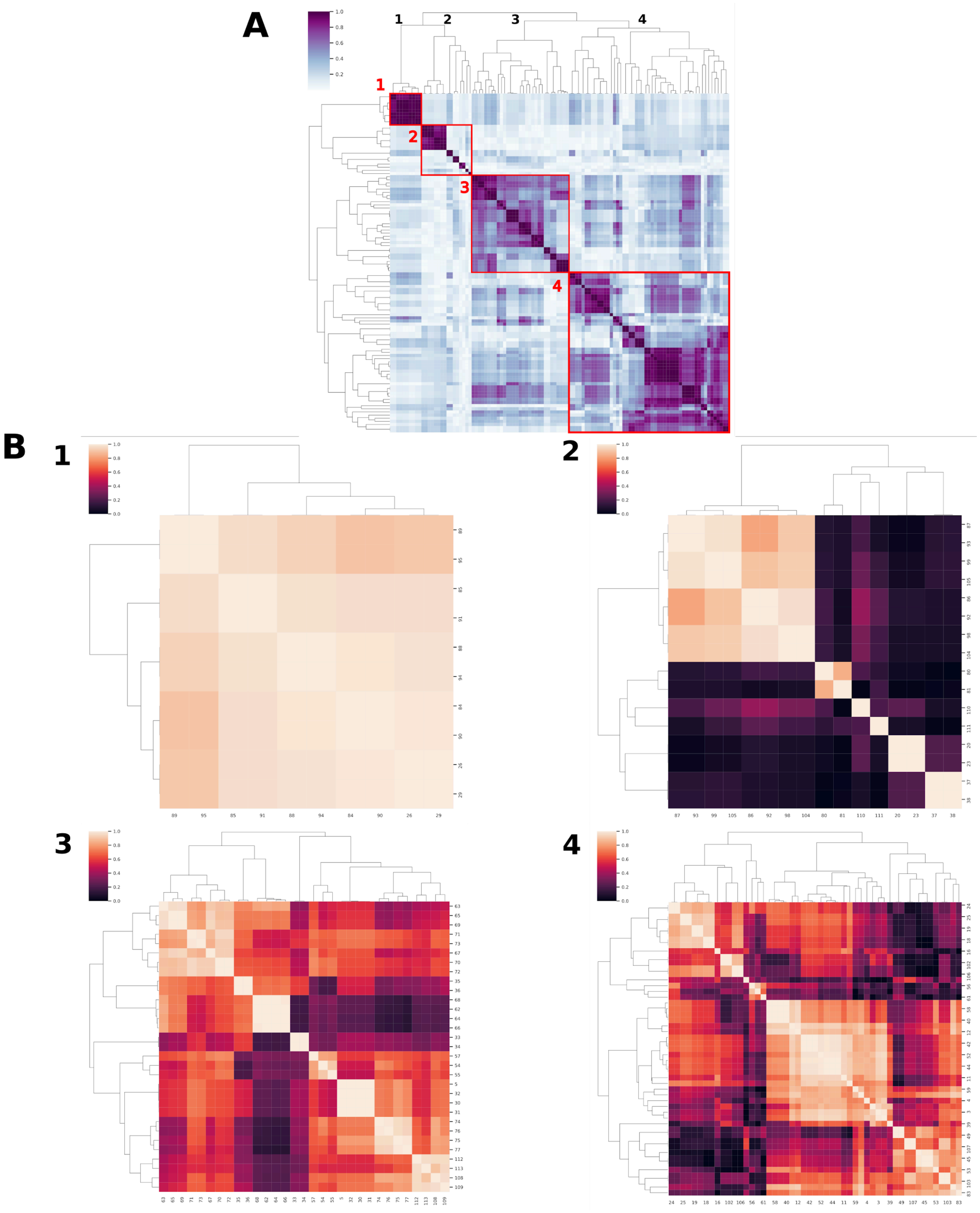


**Supplementary Figure 5**: Correlation among graph statistics.
Using various graphs constructed from a single dataset but with different parameters, we assessed the correlation among their statistics. The heatmap reveals that numerous statistics display strong correlations, suggesting they capture similar graph features. On top, the total amount of statistics is shown. The bottom plots represent the most distinct clades based on the hierarchical clustering (method: “average”).

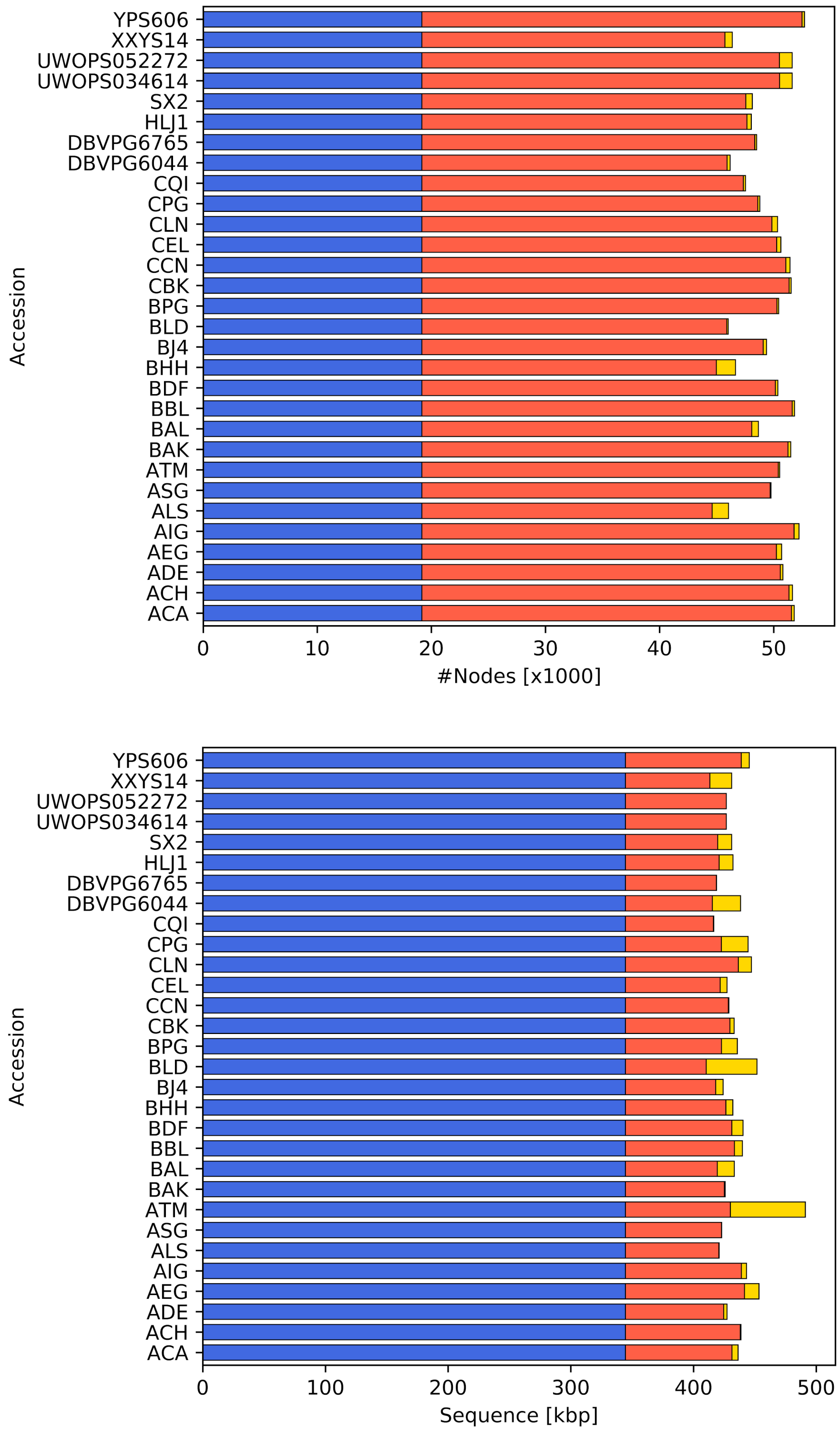


**Supplementary Figure 6:** ***gretl*** *ps* subcommand - Distribution of nodes across various *S. cerevisiae* genomes. (see Methods 1.2.4).
The top plot displays the amount of nodes in each class, the bottom plot highlights the amount of sequence for each accession. The genomes (paths) are not in any particular order.

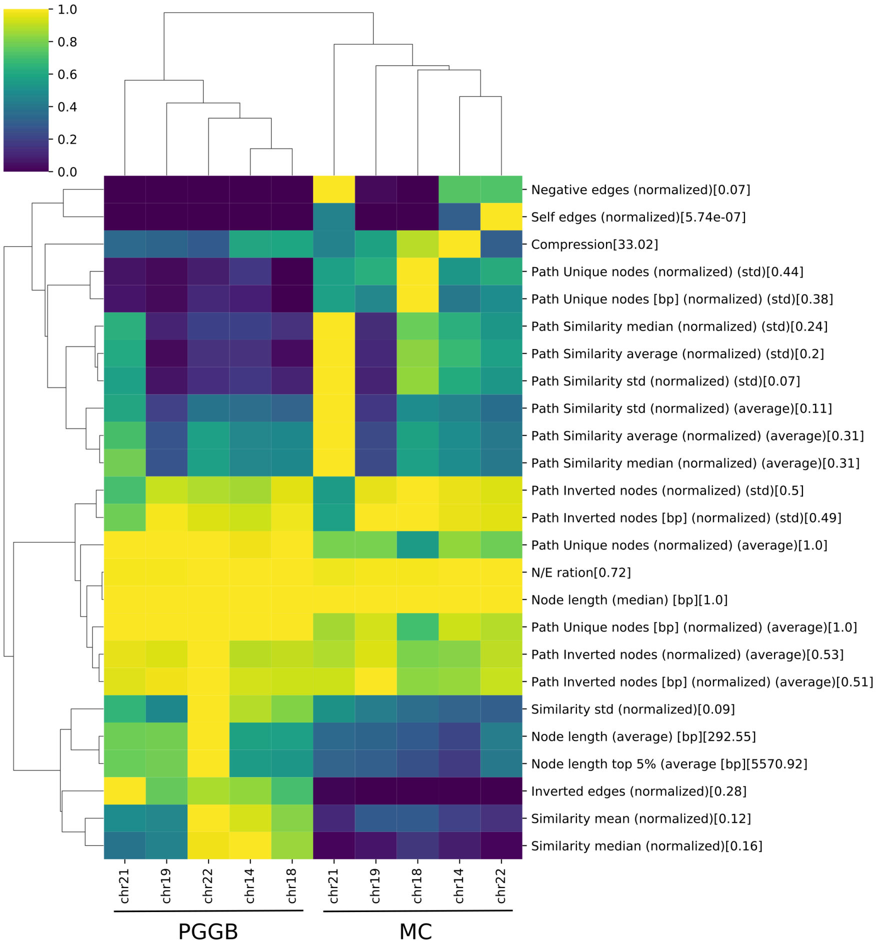


**Supplementary Figure 7: *gretl* *stats* -** Comparing different methods.
Comparing genome graphs built with different methods. Individual graphs cluster by method and not by chromosome. Only ‘normalised’ metrics are shown in this plot. In addition, values were scaled by maximum for each feature (rows). Maximum value is shown in brackets.

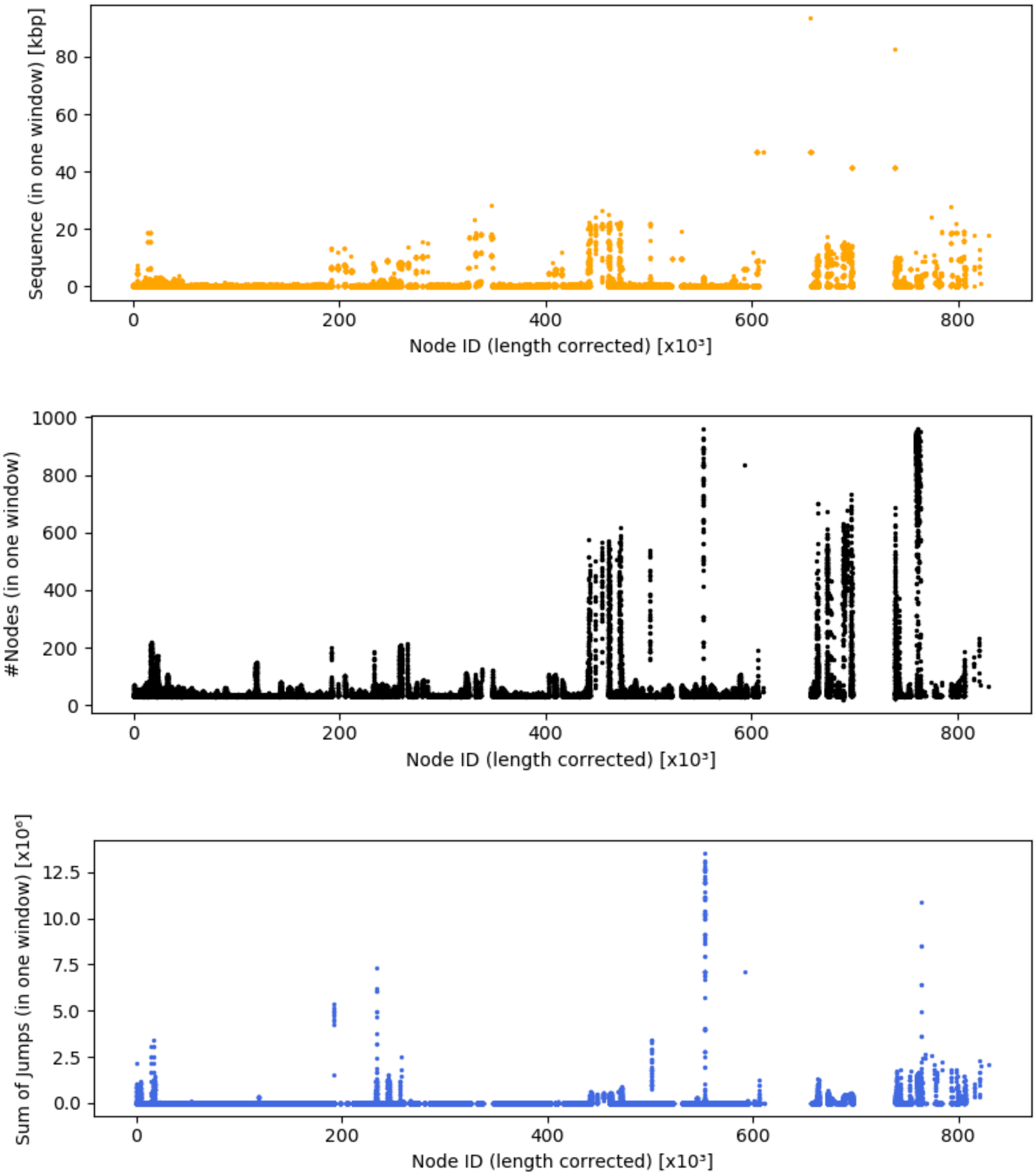


**Supplementary Figure 7:** ***gretl*** *nwindow* - Detecting regions of high local variability. Graph-based window approach iterating over each node in the graph and capturing all nodes in the range of 10 steps away. Each “window” is then summarised by (top) amount of sequence, (middle) number of nodes or (bottom) summary of node id distance (jumps) from the starting node. X-axis is inflated by node size, therefore corrected for leaving “holes” for big nodes and high frequency in arrays of small nodes.
